# The uncertainty of old aliquots of cell lines: e-CAS or m-CAS?

**DOI:** 10.1101/204149

**Authors:** Elizabeth Evans, Romain Paillot, María Rocío López-Álvarez

**Author notes:** Corresponding Author. Mailing Address: Animal Health Trust, Centre of Preventative Medicine, Lanwades Park, Newmarket, Suffolk, CB8 7UU, United Kingdom.

## Abstract

The 3Rs principles (Replacement, Reduction and Refinement) are focused on finding alternatives to the use of animals in research. In this regard, cell lines are popular and useful tools for the replacement of primary cells in *in vitro* studies. However, around 15-30% of cell lines used in research have been misidentified or cross-contaminated generating concerns about the results obtained from experiments that use them. Here we described how old aliquots of an equine macrophage cell line (e-CAS) stored at the Animal Health Trust did not contain equine cells but macrophages of murine origin (m-CAS).

## Introduction

Scientists are looking for non-sentient alternatives to support the Replacement, Reduction and Refinement (3Rs) principles and minimise the use of animal’s samples in research (https://www.nc3rs.org.uk/the-3rs). In this sense, cell lines are the most cost effective and popular replacement for primary cells thanks to their unlimited life span and ability to grow indefinitely without compromising reliable results. However, the use of cell lines comes with associated challenges like genomic variations and alteration of phenotypic characteristics of the cells over time (Maitra et al., 2005; Parson et al., 2005; Sato et al., 2016). Also, the possibility of misidentification through contamination by other cells cannot be discarded and could lead to the use of the wrong cell line (Lorsch et al., 2014). Several studies have shown that around 15-30% of cell lines used in research have been misidentified (MacLeod et al, 2008; Lacroix, 2008), creating problems and concerns about the results obtained from them. Therefore, it is essential to appropriately characterise cell lines before their use (Zhao et al., 2011; Geraghty et al., 2014; Uchio-Yamada et al., 2017).

As scientists working in equine immunology, our objective is to obtain reliable and relevant results whilst minimising the use of samples obtained from animals. Our current research is focused on the innate immune response to *Streptococcus equi (S. equi)*, a horse restricted pathogen, and how the equine innate immune cells, essentially equine macrophages and equine neutrophils, respond to S. *equi* infection *in vitro.*

The availability of equine cell lines in the market is quite limited. The ATCC only provides one horse fibroblast cell line (ATCC^®^ CCL-57™); it also offers horse tissue of unknown origin (ATCC® CRL-6583™) for which they are not responsible for its production and characterisation. Therefore, the use of equine cell lines to meet the 3Rs principles is limited and relies on aliquots kindly provided by other scientists. e-CAS is an equine macrophage cell line developed to study equine-specific pathogenic pathways and therapeutic modulation of macrophage response to defined stimuli (Werners et al., 2004). As equine macrophages, this cell line is ideal for evaluating the innate immune response to *S. equi*, and aliquots of e-CAS cells have been available at the Animal Health Trust (AHT) since 2003. Our aim was to detect variations in the expression of Toll-Like Receptors (*TLR*) in these cells by reverse transcriptase-quantitative PCR (RT-qPCR) after their stimulation with different mitogens or S. *equi* strains.

Here we describe how during the development of a PCR reaction specific for equine *TLR6* we realised that the aliquots of e-CAS cells at the AHT did not contain an equine macrophage cell line, but macrophages of murine origin.

## Materials and methods

### Cell lines

Two cell lines were used in this study. e-CAS cell line is described as equine macrophages derived from bone marrow of two 10-year-old horses (Werners et al., 2004); eqT8888 are equine cells of lymphoid origin but not completely characterised (Hormanski et al., 1992).

### DNA samples

DNA from e-CAS and eqT8888 cell lines was extracted using a QIAamp DNA minikit (QIAGEN) following manufacturer’s instructions. Archived DNA aliquots from two ponies (P1 and P2) were used as controls.

### Reactions for equine TLRs

Previously published gene specific oligonucleotide primers were used to amplify equine *TLR2* and *TLR4* (Gornik et al., 2011; Tab. 1). Specific oligonucleotide primer pairs for equine *TLR1* and *TLR6* were designed using Primer3 software (Koressaar et al., 2007; Untergasser et al., 2012), ensuring the primers completely match all the sequences available in the National Centre for Biotechnology Information (NCBI) database and did not fall within polymorphic areas of any of the genes. PCR was performed using 2μM of each primer, 1.5 μM of MgCl_2_, 200 μM dNTP, 10ng DNA template and 0.25μl Hotstar taq polymerase. Cycling conditions were: enzyme activation at 95°C for 15 min, 35 cycles of 95°C for 30 s, 55°C for 30 s and 72°C for 1 min, and a final extension at 72°C for 10 min. After the PCR process, amplified DNA fragments were resolved at 115V for 45 minutes using a 1.5% agarose gel stained with GelRed (Biotium).

**Table 1.**
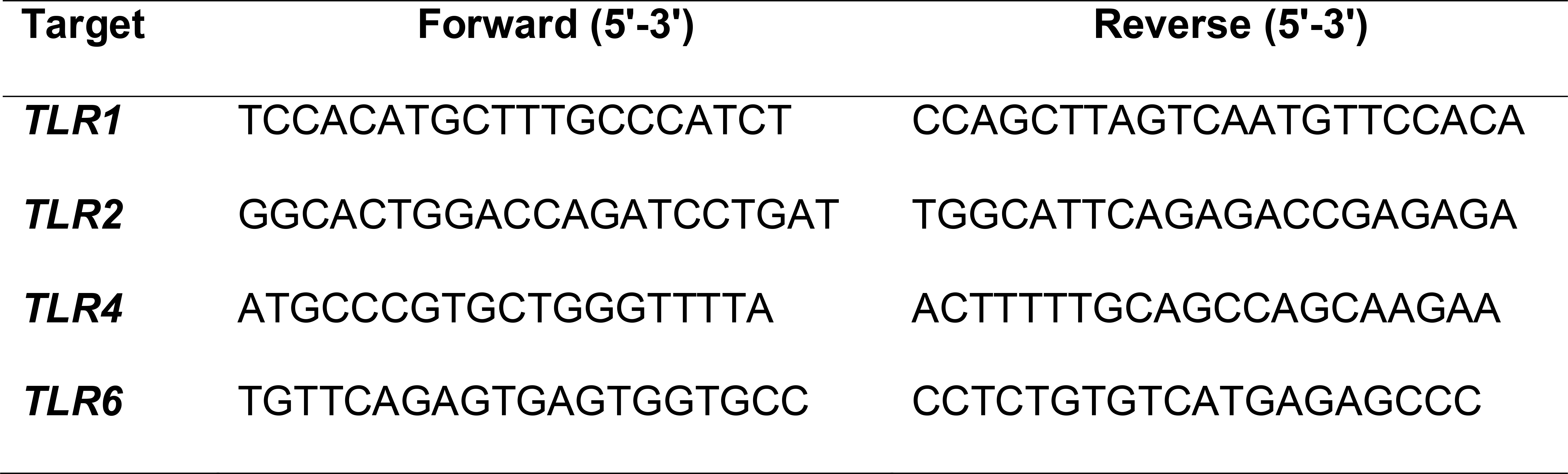
Primers used to detect equine *TLRs*

### TLR6 Sequencing

Specific amplified products from the *TLR6* PCR-SSP reaction were used for sequencing. The amplicons were clean using a Monarch® PCR & DNA Cleanup Kit (New England Biolabs) following manufacturer’s instructions. One reaction for each primer (forward and reverse) was prepared in an Axygen 96 well plate. Each reaction contained 5μl sequencing buffer, 5μl water, 1μl BigDye Terminator v3.1 (Thermofisher), 1μM forward or reverse primers and 2.5μl DNA. Cycling conditions were: 25 cycles at 96°C for 15 s, 50°C for 10 s and 60°C for 2 min, followed by 10°C indefinitely. The sequencing PCR reactions were purified by ethanol precipitation. 10μl Hi-di was then added to each well and samples were denatured in a PCR machine at 95°C for 1 min. The sequencing was performed on the ABI 3130XL genetic analyser (Applied Biosystems). Archived DNA samples from two Welsh Mountain ponies (P1, P2) were used as equine controls.

Results were analysed using Chromas (Technelysium) and aligned using Mega software (Tamura et al., 2007).

### DNA profiling

A short tandem repeat (STR) analysis was performed by the Equine Genetic Service at the AHT, an ISAG (International Society for Animal Genetics) registered laboratory. Genomic DNA samples from eCAS, eqT8888 and pony P1 were tested. The equine STR marker panel consisted of alleles at 18 different microsatellite loci: AHT4, AHT5, ASB2, ASB17, ASB23, CA425, HMS1, HMS2, HMS3, HMS6, HMS7, HTG4, HTG6, HTG7, HTG10, LEX3, LEX33 and VHL20. PCR were performed in a G-Storm GS4 Multi Block Thermal Cycler (G-storm). PCR products were separated in a capillary electrophoresis system on the 3130XL Genetic Analyser (Applied Biosystems). The GeneScan-500 LIZ Size Standard was used in each sample run for an application of automated DNA fragments analysis with five fluorescent dyes. Analysis of DNA profiles for 18 STR loci was conducted using GeneMapper 4.0 software (Applied Biosystems).

e-CAS cell line DNA profiling was performed by Mycrosynth AG (Switzerland) following the STR analysis developed by Almeida et al. (2014) for the authentication of mouse cell lines.

## Results

### Specificity of TLR reactions

The specificity of the newly developed reactions for *TLR1* and *TLR6* was determined by PCR-SSP in DNA samples obtained from e-CAS and eqT8888 cell lines. Previously developed *TLR2* and *TLR4* reactions (Gornik et al., 2011; Tab 1) were also tested alongside. As shown in Fig. 1, amplification of e-CAS DNA was only detected for *TLR6* (Fig. 1A). DNA from eqT8888 cells showed a clear band corresponding to the expected size for each of the reactions considered (Fig. 1B). The same reactions were also carried out using archived DNA samples from ponies P1 and P2, obtaining specific amplification for all *TLR* reactions (data not shown).

**Figure1.**
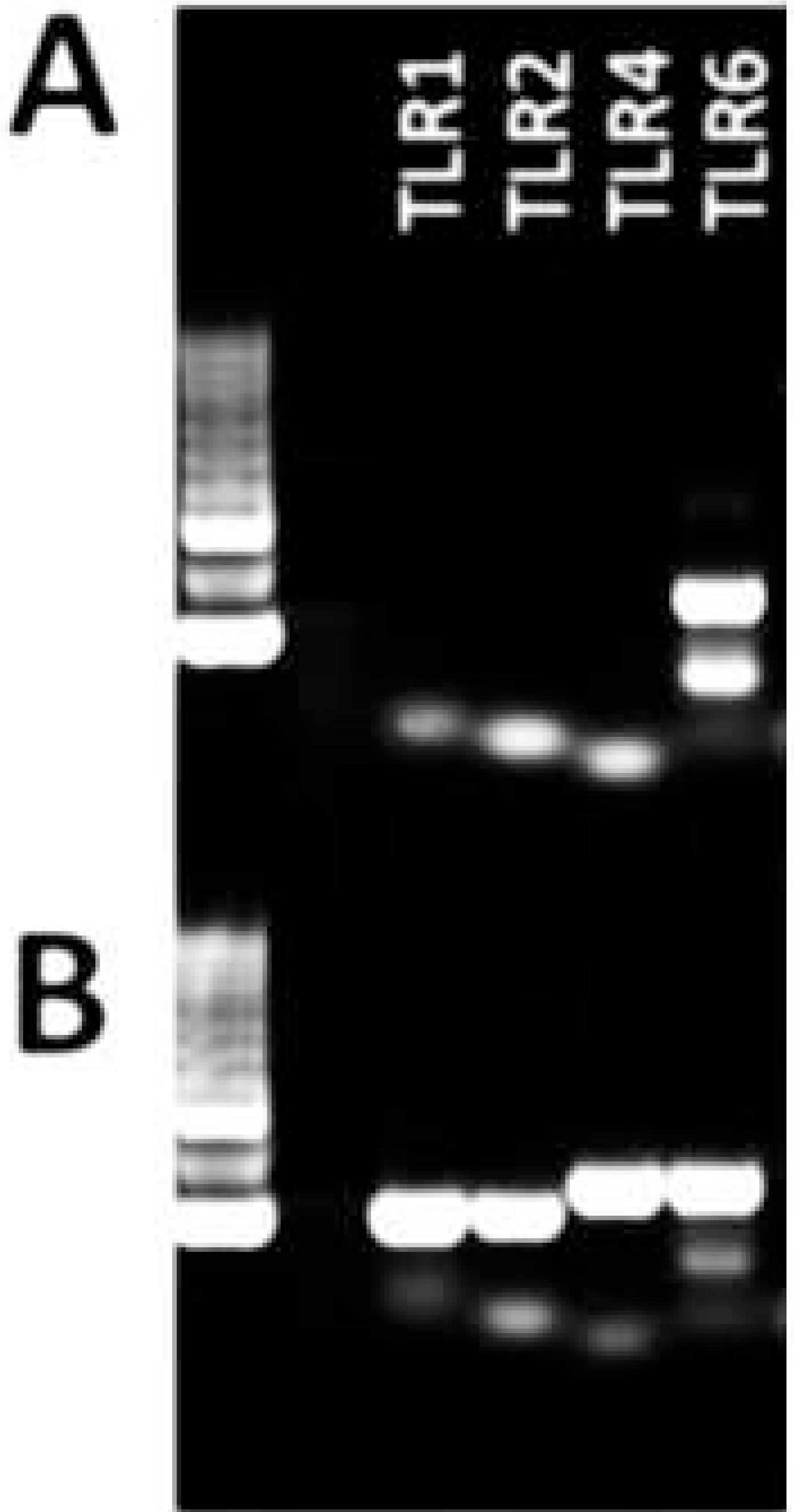
PCR reactions for equine *TLRs.* Reactions performed with A, DNA sample obtained from e-CAS cells and B, DNA sample obtained from eqT8888 cells. L: Ladder.

### TLR6 sequencing

Sequencing of the *TLR6* gene from e-CAS, eqT8888, P1 and P2 revealed up to 10 single nucleotide polymorphisms (SNPs) in the sequence obtained from e-CAS cells when compared with the results obtained from eqT8888 cells or archived DNA from ponies (Fig. 2; Supplementary Fig. S1). Sequences obtained from eqT8888, P1 and P2 DNA samples matched *TLR6* from *Equus caballus* when compared to those available in the NCBI database using Basic Local Alignment Search Tool (BLAST). However, the sequence obtained from e-CAS DNA sample shared a 99% identity with *TLR6* from *Mus musculus* (Fig. 3).

**Figure 2.**
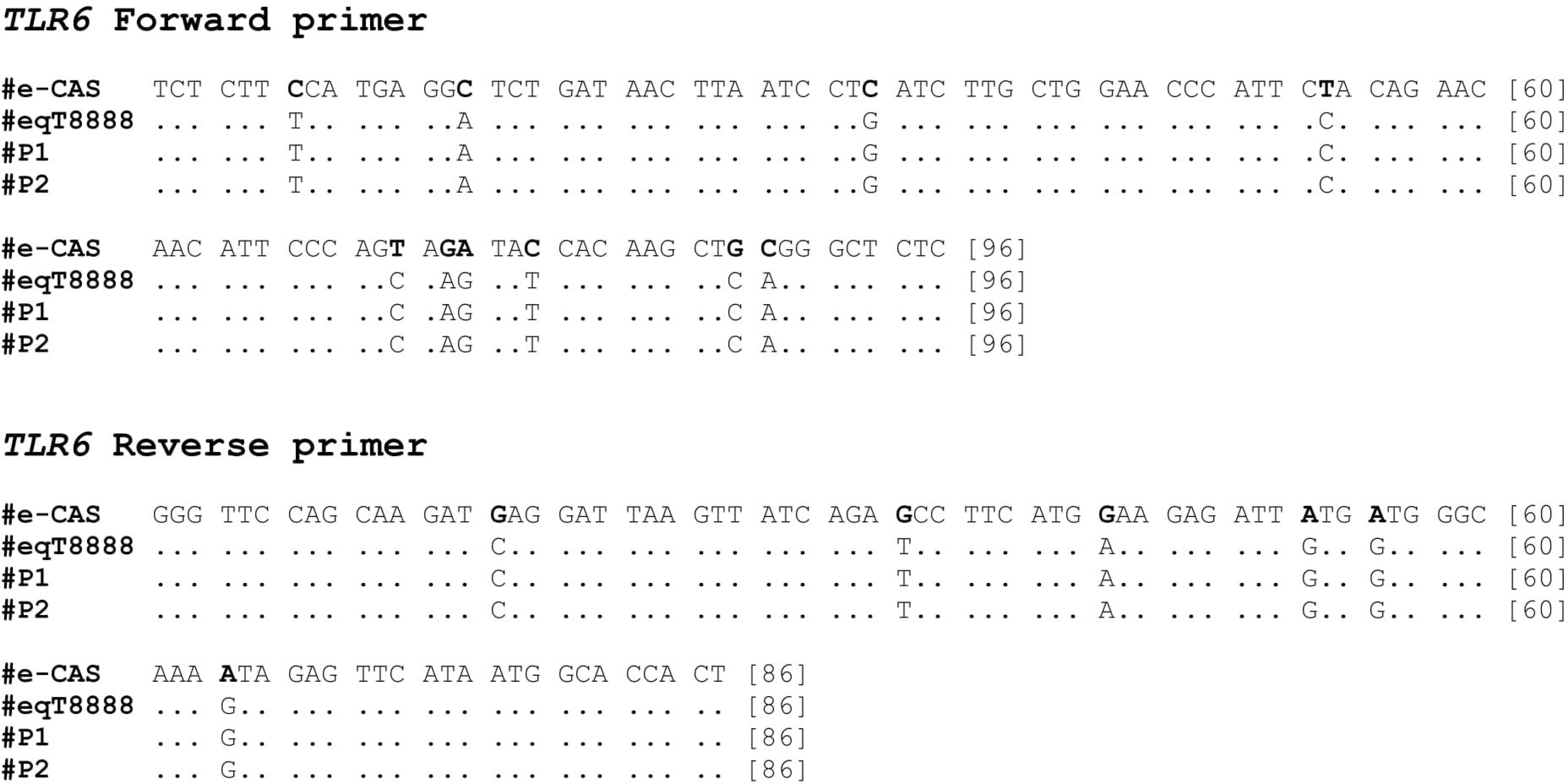
*TLR6* sequences alignment. Results obtained from sequencing of e-CAS, eqT8888, P1 and P2 DNA samples using the primers forward (top) and reverse (bottom) primers for the TLR6 reaction. SNPs are in bold.

**Figure 3.**
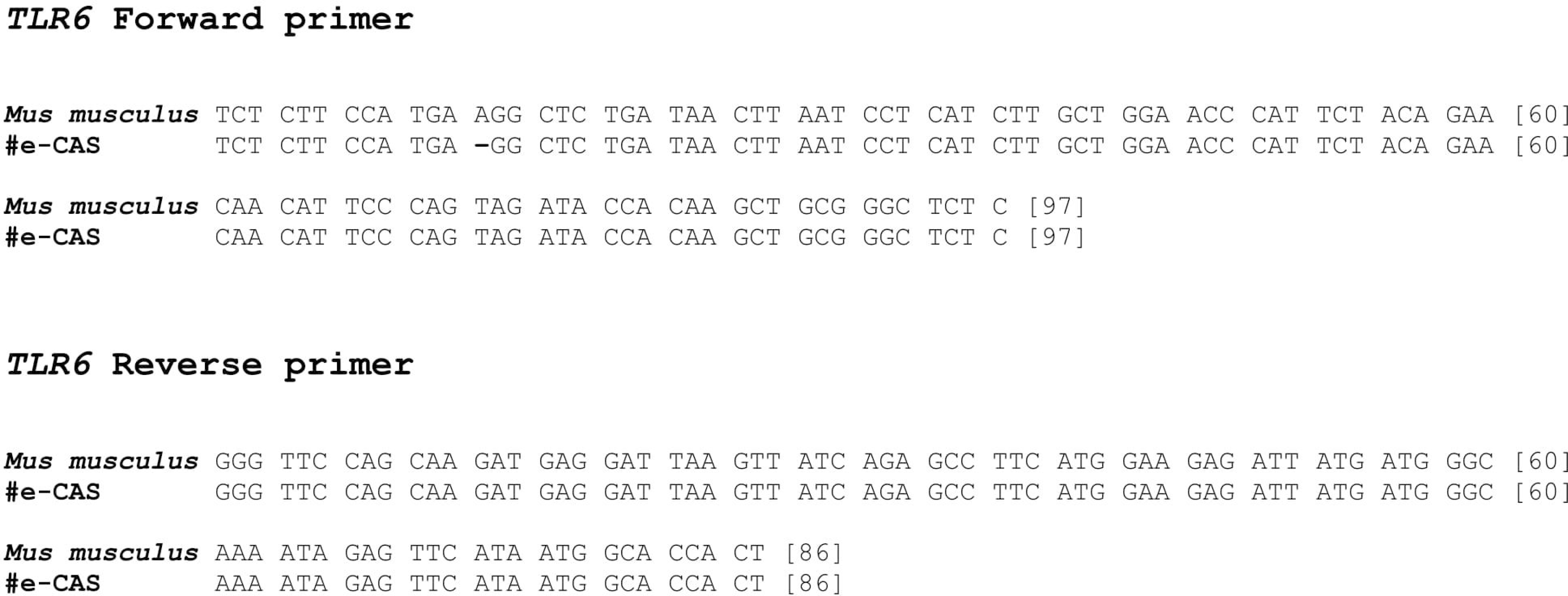
e-CAS and mouse *TLR6* sequences alignment. Results obtained from sequencing of e-CAS DNA samples were compared to the sequences available in the NCBI using BLAST. The figure shows the alignment of *TLR6* sequences from e-CAS cells and mouse.

### Short tandem repeats analysis

Short tandem repeats (STRs) are short DNA sequences of normally 2-6 bp in length that can be easily amplified by PCR. Currently, STR profiling is the standard tool for cross-contamination and authentication of human and cell lines from other species (Dirks and Drexler, 2013; Almeida et al., 2014). In order to confirm the results obtained from the *TLR6* sequencing, an equine DNA profiling by STRs analysis was performed in DNA samples from e-CAS, eqT8888 and P1. Peaks for samples from eqT8888 and P1 could be detected, analysed and alleles were assigned to each marker according to the assignment on International Society of Animal Genetics (ISAG) horse comparison test (Tab. 2). However, no results were obtained from e-CAS cells.

**Table 2:** Equine DNA profile of eqT8888 cell line and a control pony

| Alleles* |  |  |  |  |  |
| --- | --- | --- | --- | --- | --- |
| Marker | eqT8888 | Pony | Marker | eqT8888 | Pony |
| **AHT4** | H | H | **HMS6** | KM | LP |
| **AHT5** | KN | JO | **HMS7** | MN | KO |
| **ASB2** | R | KQ | **HTG4** | K | M |
| **ASB17** | GO | OR | **HTG6** | GJ | O |
| **ASB23** | KS | JU | **HTG7** | N | KO |
| **CA425** | N | J | **HTG10** | MO | O |
| **HMS1** | IJ | J | **LEX3** | H | K |
| **HMS2** | L | K | **LEX33** | M | KL |
| **HMS3** | IO | MP | **VHL20** | IN | MO |

\*according to the assignment on International Society of Animal Genetics (ISAG) horse comparison test.

Considering the results obtained for eCAS cells from the BLAST alignment and the lack of amplification during the equine DNA profiling, an aliquot of e-CAS DNA was analysed by Dr Georges Wigger (Microsynth AG, Switzerland) using the mouse DNA profiling STR test described by Almeida et al. in 2014. STRs in 9 murine loci were positively identified (Fig. 4) in eCAS DNA. The different chromosomal locations and fragment lengths are shown in Tab. 3. No trace of contamination with another mouse or human cell lines was detected.

**Figure 4.**
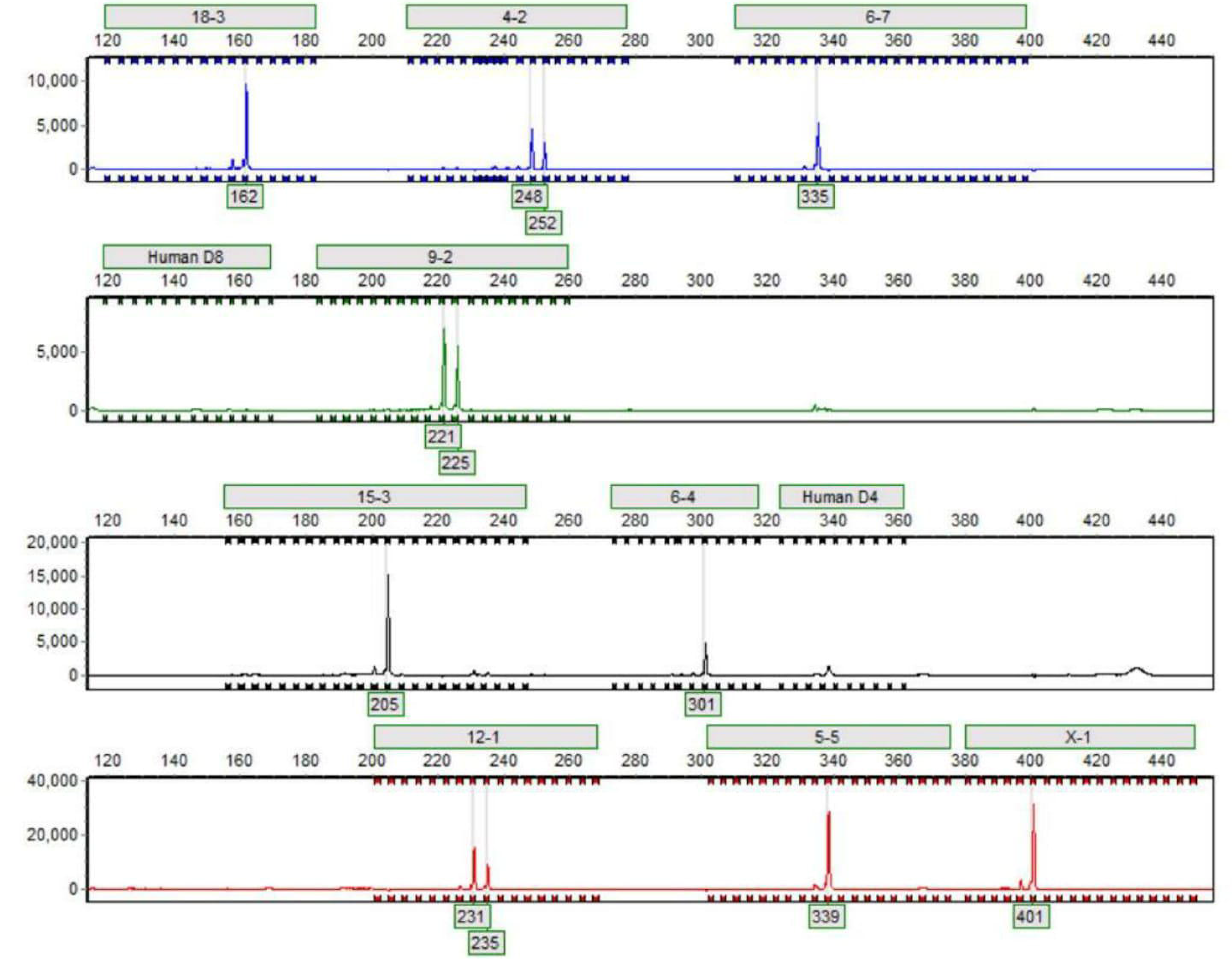
Mouse DNA profiling of e-CAS cells. The figure shows the electropherogram obtained from the DNA profiling of e-CAS cells. 9 murine markers were positively identified in the assay. Possible contamination with another mouse or human cell line was discarded.

**Table 3:** Mouse DNA profile of e-CAS cell line

| Locus | Chromosomal location | Fragment lengths |
| --- | --- | --- |
| 18-3 | Chr18 | 162 |
| 4-2 | Chr04 | 248/252 |
| 6-7 | Chr06 | 335 |
| 9-2 | Chr09 | 221/225 |
| 15-3 | Chr15 | 205 |
| 6-4 | Chr06 | 301 |
| 12-1 | Chr12 | 231/235 |
| 5-5 | Chr05 | 339 |
| X-1 | X | 401 |
| Human D8/D4 | - | - |

\*DNA profile performed by Microsynth AG using highly-polymorphic short tandem repeat loci (STR) published by Almeida et al. (2014)

## Discussion

When we conceived the project, e-CAS cells were considered as a replacement for primary macrophages obtained from ponies in order to determine variations in the expression of *TLRs* in response to different stimulatory conditions. e-CAS cells were derived from bone marrow of two 10-year-old horses to generate an equine macrophage cell line suitable for the study of equine-specific pathogenic pathways of macrophage response to defined stimuli (Werners et al., 2004) and, therefore, the ideal candidates for our work. However, DNA samples isolated from e-CAS cells stored at the AHT did not show amplification for any of the equine *TLR* reactions considered in our study except for equine *TLR6* (Fig. 1A). In contrast, specific amplification was observed for each reaction when DNA from eqT8888 cells (Fig. 1B) or samples from control ponies (data not shown) were used. The PCR reactions were designed for equine *TLRs* avoiding any known polymorphism in these genes. Amplicons were observed in 3 out of the 4 samples used, suggesting that the lack of amplification was not a technical problem of the PCR but something related to the DNA sample obtained from e-CAS cells.

The products amplified in the *TLR6* reactions for each sample were sequenced and, after aligning the sequences, it could be observed that e-CAS sample differed from eqT8888, P1 and P2 samples. Results from their comparison to sequences in the NCBI database showed that sequences obtained from eqT8888, P1 and P2 DNA samples matched *TLR6 Equus caballus*, whilst the sequence obtained from e-CAS cells shared a 99% identity with *TLR6* from *Mus musculus* (Fig. 3). These results were further confirmed by DNA profiling (Tab. 2 and Tab. 3; Fig. 4).

Cross-contamination between cell lines is quite frequent (Huang et al., 2017; Fusenig et al., 2017) and could explain the non-equine nature of e-CAS cells. During the differentiation and development of e-CAS cells, their expression of CD14 was compared with a mouse macrophage cell line (RAW264.7) to clearly establish the identity of e-CAS cells as macrophages (Werners et al., 2004). It could be plausible that the fitter and faster growing murine cell line took over the cell culture replacing the original equine macrophages. However, the DNA profile obtained from e-CAS does not exactly match the one described for RAW264.7 in 2014 by Almeida et al., discrepancy that could be explained by alterations in their fingerprint profile with prolonged culture, as has been described for other cell lines (Parson et al., 2005).

In 2007, Steinbach et al. published a study in which they evaluated the crossreactivity of anti-human leukocyte monoclonal antibodies using e-CAS and eqT8888 cell lines (Steinbach et al., 2007). The authors concluded that e-CAS cells represented an early stage of differentiation in which the characteristic macrophage surface antigens could not be detected. Nonetheless, they were clearly positive when they used an anti-mouse CD11b antibody, result that could also support the fact that the e-CAS cells used in their study were of murine origin.

To our knowledge, e-CAS cells have been used in two other publications (Wijnker et al., 2004; Lankveld et al., 2005). Wijnker et al. showed that the TNF-α release in both e-CAS cells and the human cell line U937 was similarly affected by the presence of a TNF-α converting enzyme (TACE) inhibitor (Wijnker et al., 2004). Lankveld et al. investigated the effect of ketamine on TNF-α and IL-6 release and observed that ketamine suppress LPS-induced TNF-α concentration in both e-CAS and U937 cells in a dose dependent manner and the effect of ketamine was also evident on the concentration of IL-6 released by e-CAS cells (Lankveld et al., 2005). The assays used by both groups were developed for their use in different species; they were not specific methods to measure equine cytokines (Pauli et al., 1994; Okada et al., 1997). Three other publications (Fidalgo-Carvalho et al., 2009; Karagianni et al., 2013; Swaminathan et al., 2016) have referenced Werners et al. (2004), but in all cases the citation was related to the method used during the development and differentiation of e-CAS cells, and was not associated with the characteristics of these cells as equine macrophages. To our knowledge, there is no other publication in which e-CAS cells have been used in equine-specific experiments.

In summary, this work shows that the aliquots of e-CAS cells stored at the AHT are of murine origin rather than equine macrophages. However, this established identity was only possible after performing molecular assays that are specific for the equine species.

In conclusion, the data presented here highlight the importance of cell line authentication prior to their use in research.

## Acknowledgements

The authors would like to thank Claire Trushell and Angie Stagg from the AHT Equine Genetic Service for kindly performing the STR analysis of our samples. This work was funded by the Pet Plan Charitable Trust (PPCT 204-278) and by the Animal Welfare Foundation through the Norman Hayward Research Fund (NHF_2016_02_MLA).

